# Improved Genome Assembly of *Onygena corvina*, a Keratin-Degrading Fungus

**DOI:** 10.1101/2025.07.02.662733

**Authors:** Siddhi Pavale, Vincent G.H. Eijsink, Sabina Leanti La Rosa

## Abstract

**Objective:** *Onygena corvina* is a non-pathogenic, saprophytic fungus that colonizes feathers, hooves, and hair, and represents a valuable source of keratin-degrading enzymes. Previously, the genome of *O. corvina* has been assembled based on Illumina short-read sequencing, yielding a reference genome composed of 521 contigs with a contig N50 of 0.229 Mb.

**Results:** Here, we report an improved *O. corvina* genome assembly generated using a high-quality hybrid approach that combines Illumina short-read and Oxford Nanopore long-read sequencing. The new assembly consists of only 13 contigs totaling 21.8 Mb, with an N50 of 4.4 Mb, and has a completeness of 98.97%. A total of 7,232 protein-coding genes were identified using an integrative approach that combines *de novo* predictions, homology-based inferences, and RNA-sequencing–guided evidence. Notably, 158 putative protease-coding genes were identified representing a substantial increase from the 73 predicted proteases in the previous annotation. Our improved genome assembly and associated gene annotations will facilitate comparative genomics, and high-resolution mapping of transcriptomic and proteomic data, to advance research on fungal physiology and fungal abilities to degrade recalcitrant substrates such as keratin.

## Introduction

*Onygena corvina* is a saprotrophic ascomycete in the order Onygenales, capable of colonizing keratinous materials such as feathers, hooves, and hair [1, 2]. Its keratinolytic potential, coupled with a non-pathogenic nature, makes this fungus a promising candidate for keratin waste valorization [3, 4]. However, the existing reference genome is fragmented and only partially annotated, limiting in-depth functional analyses [5, 6]. Here, we present a high-quality genome assembly for *O. corvina* and associated gene annotations, which were obtained using a hybrid approach combining Illumina short-read and Oxford Nanopore long-read sequencing for DNA as well as Illumina sequencing for RNA.

## Main Text

### Genome Assembly and Annotation

*O. corvina* CBS 281.48 was obtained from the CBS-KNAW Collection of the Westerdijk Fungal Biodiversity Institute (Utrecht, Netherlands). For DNA and RNA extraction, mycelial biomass was generated in 500 ml flasks with 200 ml potato dextrose broth that were inoculated with five to eight mycelial plugs and incubated at 25 °C, 100 rpm for 7 days. Mycelia were harvested using miracoth filters (Merck KGaA, Darmstadt, Germany), washed with distilled water, flash-frozen in liquid nitrogen, and stored at – 80 °C until further use.

Genomic DNA was extracted using the ZYMO Research Quick-DNA Miniprep Plus Kit (Zymo Research, Freiburg, Germany). For long-read sequencing, a library was prepared with a Native Barcoding Expansion Kit (EXP-NBD104, Oxford Nanopore Technologies, Oxford, UK) and sequenced on a Nanopore PromethION 48 with a Flow Cell (R10.4.1, FLO-PRO114M). Library construction and sequencing were performed at Biomarker Technologies GmbH (Münster, Germany). Base-calling and adapter trimming were performed using Dorado v0.7.2 (Oxford Nanopore Technologies) in super-accurate mode. The long-read sequencing generated 380,647 reads, totaling approximately 2.33 Gb, with an average length of 6,114 bp and with an N50 of 7,952 bp. For short-read sequencing, a sample library was prepared for Illumina pair-end sequencing (2 × 150 bp) with the Nextera DNA Flex kit (Illumina, San Diego, USA) followed by sequencing with an Illumina NovaSeq PE150 platform. Overall, the sequencing generated an estimated genome coverage of 106 X.

Total RNA was extracted using the ZYMO Research Quick-RNA Miniprep Plus Kit following the manufacturer’s instructions. The RNA-seq library was constructed using the Ultima Dual-mode mRNA Library Prep Kit for Illumina (Yeasen Biotechnology, Shanghai, China) following the manufacturer’s instructions. The library was then sequenced on the Illumina NovaSeq X platform, yielding 6.77 Gb of 150-bp paired-end reads.

A hybrid *de novo* assembly of the short and long reads was obtained using Canu [7], wtdbg [8], and Racon [9]. A final round of error correction using Pilon [10] and short-read Illumina data ensured high base-level accuracy. There were no assembly gaps, and quality metrics indicated 99.96% coverage with an average sequencing depth of 66.1X. Assessment of assembly completeness using BUSCO [11] and the fungi_odb9 dataset identified 287 of 290 (98.97%) conserved fungal genes, indicating a near-complete assembly. Repetitive sequences were identified using a species-specific repeat database constructed with LTR_FINDER [12], MITE-Hunter [13], RepeatScout [14], and PILER-DF [15]. This database was classified with PASTEClassifier [16], merged with Repbase [17], and used for annotation with RepeatMasker [18]. The analysis revealed that 3.22% of the genome consisted of repeats and that simple sequence repeats and unclassified elements made up the bulk of repetitive DNA.

Prediction of protein-coding genes was carried out by integrating *de novo*, homology-based, and transcriptome-supported approaches. Genscan [21], Augustus [22], GlimmerHMM [23], GeneID [24], and SNAP [25] were used for *de novo* prediction, while GeMoMa [26] was used to leverage homology with closely related fungal genomes (*Coccidioides immitis, Coccidioides posadasii* and *Ophidiomyces ophidiicola*). Transcriptomic data were mapped with HISAT2 [27] and assembled with StringTie [27], after which coding regions were identified using TransDecoder [28] and PASA [29]. The final gene set was constructed using EVM [30] and refined using PASA [29], resulting in 7,232 protein-coding genes. The average gene length was 2164 bp, with 3.37 exons per gene and an average coding sequence length of 479 bp. Of note, 99.01% of gene predictions were supported by transcriptome or homology evidence, indicating high reliability of these predictions.

### Comparative Genomic Analysis

Compared to the previous genome assembly (GCA_000812245.1), the updated *O. corvina* genome is more contiguous. While the total assembly length remains ∼21.8 Mb, the number of contigs was reduced from 521 to 13. The contig N50 was increased substantially from 0.229 Mb to 4.41 Mb, with the largest contig now spanning 6.85 Mb (Table 1). Sequence alignment using Minimap2 [19] and DGENIES [20] revealed 98.65 % identity with the earlier version of the genome, indicating high identity and collinearity (Figure 1).

**Table 1.** Assembly statistics for the previous (PRJNA270018) and new (PRJNA1280007) version of the *O. corvina* genome.

| Characteristic | Previous version<br>(PRJNA270018) | New version<br>(PRJNA1280007) |
| --- | --- | --- |
| Genome Size (Mb) | 21.7 | 21.8 |
| Number of contigs | 521 | 13 |
| Largest Contig size (Mb) | 0.933 | 6.854 |
| Contig N50 (Mb) | 0.229 | 4.410 |
| GC percent | 48 | 48 |

**Figure 1.**
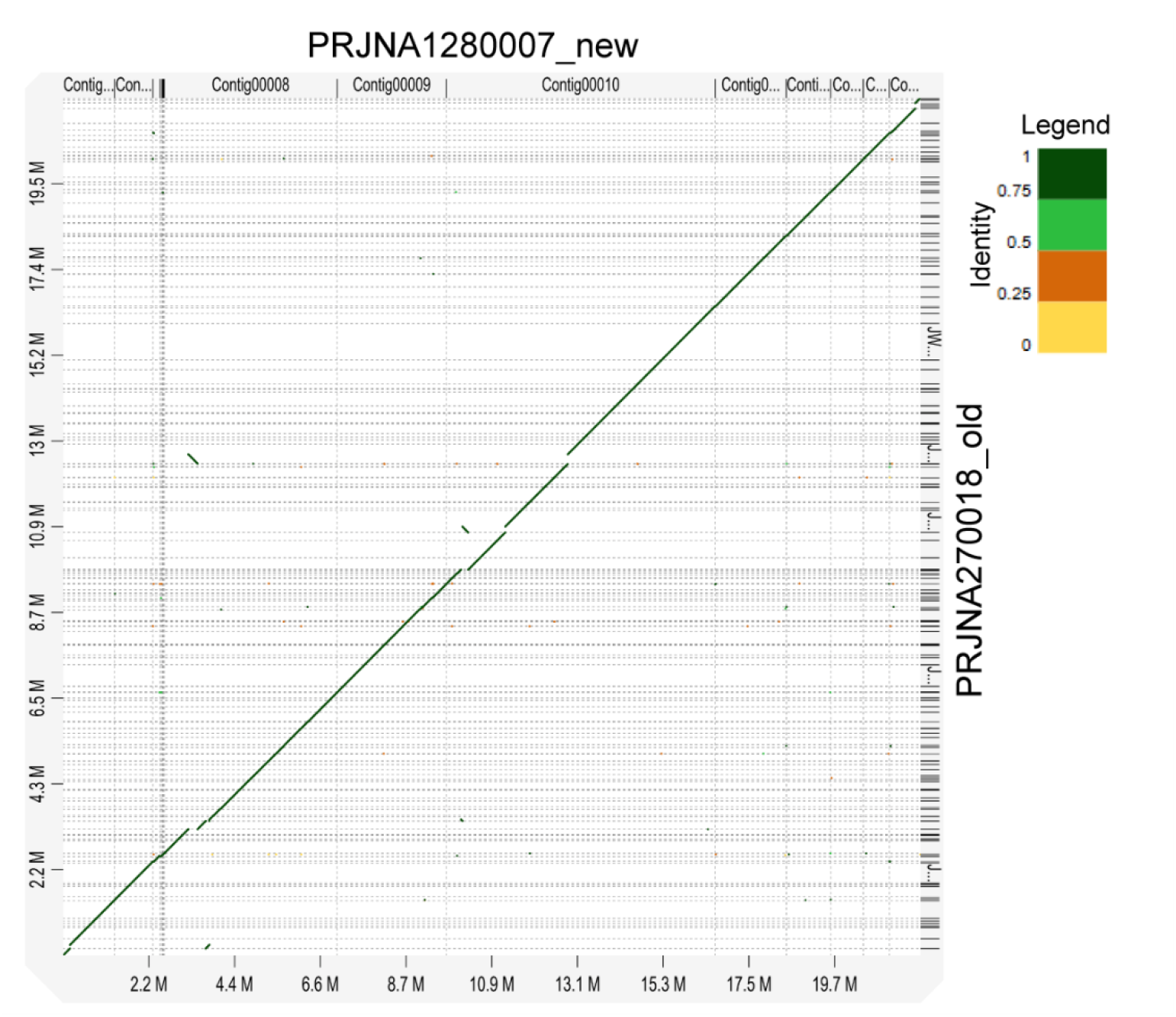
Alignment between the new version (PRJNA1280007) (top) and the previous version (PRJNA270018/GCA_000812245.1) (right) of the *O. corvina* genome. The alignment was visualized with D-GENIES, where the coloration corresponds to the percent identity as per the legend.

### Functional Genomics and Keratin Degradation Potential

We additionally conducted a secondary metabolite cluster analysis using antiSMASH (v8.0) for fungi, with the detection strictness set to relaxed [31]. A total of 32 biosynthetic gene clusters (BGCs) were identified, including terpene, type I polyketide synthase (T1PKS), non-ribosomal peptide synthetase (NRPS), and hybrid PKS-NRPS clusters. Several BGCs showed high similarity to known clusters: YWA1 [32] (polyketide); AbT1 [33] and chrysogine [34] (both NRPS); clavaric acid [35] (terpene); and tolypyridone C [36] (hybrid PKS-NRPS).

Given the ability of *Onygena corvina* to colonize and degrade keratin-rich substrates, we explored its keratinolytic potential through genome-wide prediction of proteases. Through motif recognition using the Homology to Peptide Pattern (Hotpep) pipeline, 158 proteases were predicted, a substantial expansion from the 73 identified in the previous assembly [1]. These spanned across six MEROPS [37] families including aspartic proteases (A; n=6), asparagine/peptidyl-lyases (N; n=1), metalloproteases (M; n=51), cysteine proteases (C; n=34), serine proteases (S; n=53), and threonine proteases (T; n=13). Out of 158 proteases, 59 were predicted to be secreted based on signal peptide and transmembrane domain analysis using SignalP 5.0 [38] and TMHMM [39] v2.0 (Figure 2A). Further classification based on subcellular localization showed that the secreted proteases were predominantly serine and metalloproteases, while the non-secreted proteases largely belonged to cysteine and metalloprotease families (Figure 2B). Additionally, we identified 189 carbohydrate-active enzymes (CAZymes) in the *O. corvina* genome, using dbCAN3 [40]. Of these, 38 were predicted to be secreted, including glycoside hydrolases (GH; n=27), carbohydrate esterases (CE; n=2), auxiliary activity enzymes (AA; n=8), and glycosyltransferases (GT; n=1). Of note, two genes (*Onygena_corvina0G042260.1* and *Onygena_corvina0G066340.1*) coding for lytic polysaccharide monooxygenases (LPMOs) in AA11 family were detected, corroborating earlier findings for *O. corvina* and related keratin-degrading fungi [41]. While AA11-type LPMOs are known to act on crystalline chitin [42, 43] and soluble chitin fragments [44], these redox enzymes have been proposed to facilitate cleavage of keratin, possibly by targeting its carbohydrate components [2].

**Figure 2.**
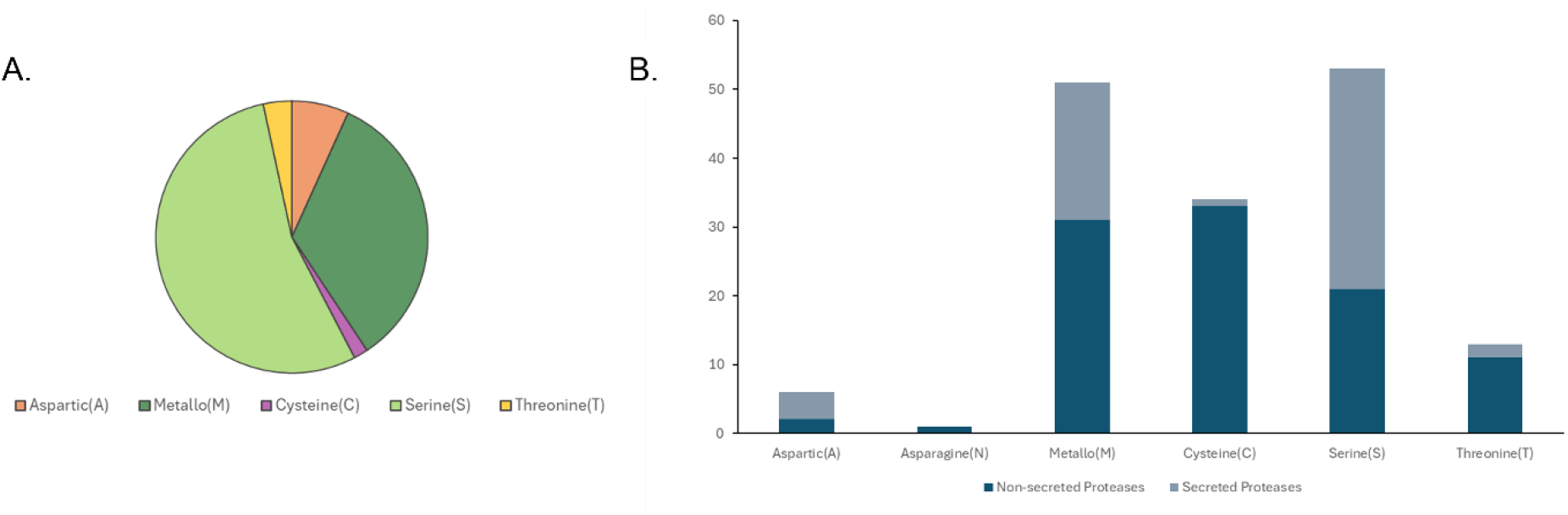
MEROPS-based classification of proteases encoded in the *O. corvina* genome, categorized into serine (S), metalloprotease (M), threonine (T), cysteine (C), asparagine (N) and aspartic (A) families. **(A)** Distribution of the 59 putatively secreted proteases across these families. **(B)** Bar chart showing the comparative abundance of secreted versus non-secreted (intracellular and membrane-bound) proteases (158 enzymes in total). Light blue bars represent secreted proteases, while dark blue bars indicate non-secreted proteases.

## Limitations

Comparative genomic analyses beyond basic assembly comparisons were not performed, and further investigations are needed to determine the phylogenetic placement and taxonomic relatedness of *O. corvina* to other keratin-degrading fungi. Additionally, functional validation of the predicted proteases and CAZymes remains to be carried out.

## Abbreviations

AA: Auxiliary activity enzymes
CAZymes: Carbohydrate-active enzymes
CE: Carbohydrate Esterases
CBS: Centraalbureau voor Schimmelcultures (Central Bureau of Fungal Cultures)
EVM: Evidence Modeler
GH: Glycoside hydrolases
GH: Glycosyltransferases
Hotpep: Homology to Peptide Pattern pipeline
LPMOs: Lytic polysaccharide monooxygenases
N50: Assembly metric indicating contig length at 50% genome coverage
PE150: Paired-end 150 base pair sequencing
PASA: Program to Assemble Spliced Alignments
SignalP: Signal peptide prediction software
TMHMM: Transmembrane helix prediction software
BUSCO: Benchmarking Universal Single-Copy Orthologs

## Declarations

- Ethics approval and consent to participate: Not Applicable
- Consent for publication: Not Applicable
- Availability of data and materials: The genome, DNA and RNA reads have been made available in the NCBI GenBank under the accession number PRJNA1280007. The gene and protein annotations, putative functional annotation, and CAZyme predictions are available on Figshare (https://doi.org/10.6084/m9.figshare.29328893.v1).
- Competing interests: Not Applicable
- Funding: This work was supported by the SFI Industrial Biotechnology program (project number 309558) funded by the Research Council of Norway.
- Authors’ contributions: SP and SLLR conceptualized the study, performed data analysis, and drafted the manuscript. SLLR and VE critically revised and reviewed the manuscript.

## Additional File

- File name: Additional File 1
- File format: Excel file (xlsx)
- Title of data: List of 38 putatively secreted carbohydrate-active enzymes encoded in the *Onygena corvina* genome.
- Description of data: For each gene, the table provides the Gene ID, predicted CAZy family categorized as glycoside hydrolases (GH), glycosyltransferases (GT), carbohydrate esterases (CE), or auxiliary activity (AA) enzymes, annotated using dbCAN3, along with known enzymatic activities in each family according to the CAZy database.

